# PI-Plat: A high-resolution image-based 3D reconstruction method to estimate growth dynamics of rice inflorescence traits

**DOI:** 10.1101/835306

**Authors:** Jaspreet Sandhu, Feiyu Zhu, Puneet Paul, Tian Gao, Balpreet K. Dhatt, Yufeng Ge, Paul Staswick, Hongfeng Yu, Harkamal Walia

## Abstract

**Background:** Recent advances in image-based plant phenotyping have improved our capability to study vegetative stage growth dynamics. However, more complex agronomic traits such as inflorescence architecture (IA), which predominantly contributes to grain crop yield are more challenging to quantify and hence are relatively less explored. Previous efforts to estimate inflorescence-related traits using image-based phenotyping have been limited to destructive end-point measurements. Development of non-destructive inflorescence phenotyping platforms could accelerate the discovery of the phenotypic variation with respect to inflorescence dynamics and mapping of the underlying genes regulating critical yield components.

**Results:** The major objective of this study is to evaluate post-fertilization development and growth dynamics of inflorescence at high spatial and temporal resolution in rice. For this, we developed the *P*anicle *I*maging *Plat*form (PI-Plat) to comprehend multi-dimensional features of IA in a non-destructive manner. We used 11 rice genotypes to capture multi-view images of primary panicle on weekly basis after the fertilization. These images were used to reconstruct a 3D point cloud of the panicle, which enabled us to extract digital traits such as voxel count and color intensity. We found that the voxel count of developing panicles is positively correlated with seed number and weight at maturity. The voxel count from developing panicles projected overall volumes that increased during the grain filling phase, wherein quantification of color intensity estimated the rate of panicle maturation. Our 3D based phenotyping solution showed superior performance compared to conventional 2D based approaches.

**Conclusions:** For harnessing the potential of the existing genetic resources, we need a comprehensive understanding of the genotype-to-phenotype relationship. Relatively low-cost sequencing platforms have facilitated high-throughput genotyping, while phenotyping, especially for complex traits, has posed major challenges for crop improvement. PI-Plat offers a low cost and high-resolution platform to phenotype inflorescence-related traits using 3D reconstruction-based approach. Further, the non-destructive nature of the platform facilitates analyses of the same panicle at multiple developmental time points, which can be utilized to explore the genetic variation for dynamic inflorescence traits in cereals.

## Background

With increasing world population, climatic variability and declining arable land resources, the need to increase global food production is paramount [1–3]. Two components that are essential for achieving global food security involve precise agronomic management and genetic improvement of major crops such as rice, wheat, and maize. Integral to both components is the development of data-driven tools that increase precision in implementation and enhance predictive capabilities. Moreover, strategic selection and adaptation of yield-related traits to maximize agricultural production holds the key to achieve sustainable food security [4–6]. Inflorescence architecture (IA) is an important phenotypic feature that ultimately contributes to most of the grain crop yield components such as grain number, size, and weight [7–9]. However, the complexity of IA, especially in cereals, is a limiting factor for accurate determination of yield traits. Estimating the yield-related traits by conventional methods is subjective, laborious, and error-prone [10]. Also, the scope of the detectable yield-related traits is limited by manual measurements, which increases the chances of damaging the inflorescence.

Advances in automation of plant phenotyping technologies, mainly in reference to image-based phenotyping, have increased the depth and the scale of measuring vegetative traits [11–19]. However, only a few studies have used the phenotyping platform to screen IA [16, 20–22]. Some platforms have utilized machine-vision-based approaches to estimate inflorescence-related parameters [23–26]. In addition, two-dimensional (2D) imaging platforms have been employed, for example, *T*assel *I*mage-based *P*henotyping *S*ystem (TIPS) quantifies morphological traits from freshly harvested maize tassels, while *PA*nicle *ST*ructure *A*nalyzer for *R*ice (PASTAR/PASTA), *P*anicle *TRA*it *P*henotyping (P-TRAP), and PANorma analyze rice panicle length and branching [20, 21, 27, 28]. Both P-TRAP and PANorma have been used for genome-wide association studies (GWAS) with respect to rice panicle traits [27, 29–31]. Recently, Zhou *et al* [22] developed *T*oolkit for *I*nflorescence *M*easurement (TIM) to estimate sorghum panicle volume derived from two planar imaging data. The derived panicle-related traits of sorghum were used for GWAS to facilitate gene discovery.

Most of these 2D image-based IA approaches have discussed only the mature or end-point traits and do not capture the growth dynamics of developing inflorescence. Furthermore, biplanar images can only provide 2D projections of a 3D structure, thus accounting for substantial loss of spatial information [32]. 3D imaging has started to gain momentum to circumvent limitations of 2D imaging [33]. Different 3D imaging methods, for example time of flight (ToF), laser scanning, stereovision among others, have been applied for remote sensing or field-based phenotyping platforms In addition, depth cameras are also widely used for capturing an entire plant or large plants parts [34]. Stereovision, which considers object images from different angles to reconstruct 3D surfaces, offers an inexpensive, accurate and efficient method for on-site 3D plant imaging [32, 35, 36]. The recent introduction of freely available software – Multi-View Environment (MVE) offers an end-to-end 3D reconstruction solution [37]. MVE combines the multi-view stereo (MVS) and structure-from-motion (SfM) algorithms to generate dense point clouds for 3D object reconstruction [37]. The MVS-SfM approach has been used to reconstruct 3D meshes of leaves, canopy or whole plant [38–41]. However, this approach has not been used to characterize IA. Here, we present the results from characterizing rice panicles using the 3D reconstruction-based approach. The main objectives of our study were to (a) capture multi-dimensional, high-resolution images of ‘panicle on plant’ after the fertilization to reconstruct 3D plant cloud of inflorescence, (b) use 3D point clouds to derive inflorescence-related traits, and (c) use the derived traits to monitor growth dynamics of developing inflorescence and distinguish inherent genetic and morphological diversity in crop species.

However, it is challenging to perform 3D reconstruction of rice panicles to achieve our objectives. First, a rice panicle is often occluded by other plant components such as leaves and other panicles. Therefore, the existing solutions by moving cameras [42] are not entirely suitable to generate un-occluded images for a panicle. Second, a panicle is non-rigid and typically is not located in the center of a plant, making it difficult to apply the existing solutions based on plant rotation [42]. Third, rather than destructive methods [22], non-destructive methods are needed to keep a panicle alive, as the growth dynamics of a panicle is of interest in this study. Fourth, the size of a panicle is relatively marginal, and the depth-camera based solutions [34] may not provide sufficient resolutions to capture the 3D details of a panicle.

To address these challenges, we developed an *in-house P*anicle *I*maging *Plat*form (PI-Plat) to capture the dynamics of developing panicles in rice from a range of genetically diverse rice genotypes. A panicle is isolated to generate un-occluded images in a non-destructive manner. In addition, a panicle stays still at the center in the PI-Plat and cameras rotate around it, thus minimizing the vibration and allowing generation of a more stable 3D point cloud. The resolution of the cameras is ensured to capture details of a panicle in 2D images, leading to high-resolution 3D reconstruction results. A total of 11 genotypes, *indica* and *japonica* sub-populations were selected. Post fertilization, primary panicles were imaged on a weekly basis (week 1, 2, and 3) by using the PI-Plat. The captured images were used for 3D reconstruction to extract digital phenotypic attributes: voxel count and color intensity. We reported increased sensitivity in panicle trait prediction from 3D reconstruction when compared to direct end-point measurements of yield components. Although the PI-Plat is designed for rice panicles, it can be extended for other small plant components such as new branches or leaves for cereals.

## Material and methods

### Plant material

Surface-sterilized seeds of 11 rice accessions were germinated on half strength Murashige and Skoog media for 3 days in dark, followed by a day in light (list of the genotypes used in the study; Additional File 1). Initially, two uniformly germinated seedlings were transplanted to a 4-inch square shaped pot filled with pasteurized field soil. Throughout the growing season, the pots were maintained in standing water. After 10 days of transplanting, seedlings were thinned to retain one plant per pot per genotype.

### Temperature treatment

Plants were grown under control conditions (16-hour light and 8-hour dark at 28±1°C and 23±1°C) till anthesis. One day after 50% of the primary panicle completely fertilized, half of the plants from each genotype were transferred to greenhouse having high night-time temperature (HNT; 16-hour light and 8-hour dark at 28±1°C and 28±1°C). HNT treatment was maintained until maturity. Two or three replicates per treatment per genotype from the current set were used to establish image-based phenotyping workflow (Figure 1).

**Figure 1:**
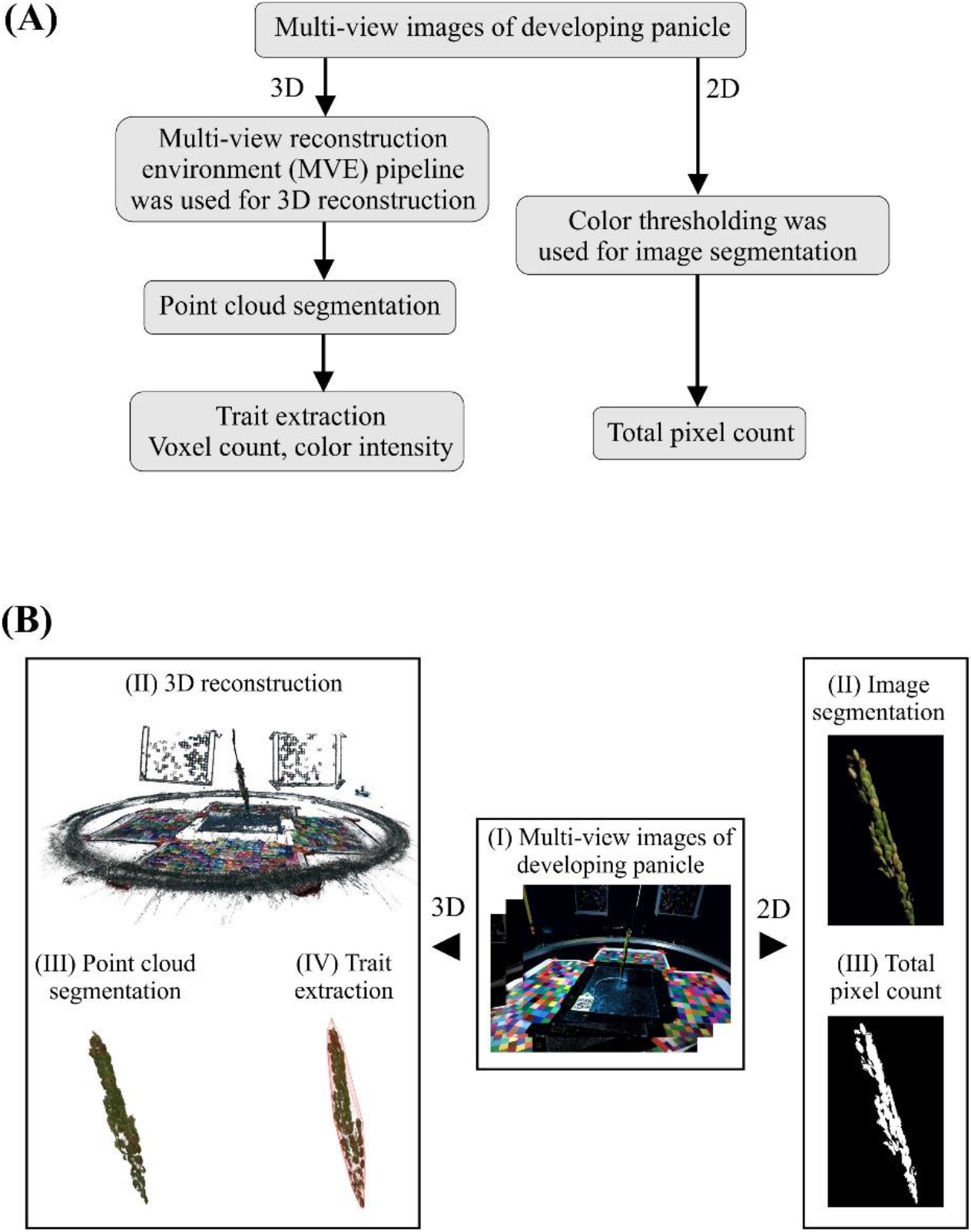
Multi-view image analysis of developing panicle using PI-Plat. **(A)** Flowchart and **(B)** graphical representation of the multi-view image analysis using 3D reconstruction and 2D approach.

### PI-Plat: Panicle Imaging Platform

We constructed a low-cost *P*anicle *I*maging *P*latform (PI-Plat) to capture the growth parameters of rice panicles after flowering (Additional File 2). The PI-Plat is comprised of three main parts: (i) a customized wooden chamber with black interior, (ii) a rotating imaging system, and (iii) color checkerboards.

#### Customized wooden chamber and rotating imaging system

To host the PI-Plat, a wooden chamber (height: 75-inch, width: 52.5-inch, length: 55-inch) was customized (Additional File 2). The interior of the chamber was painted black to reduce the light interference and increase the quality of image segmentation during the image processing procedure. Inside the chamber, a circular wooden board (diameter: 37-inch) having an aperture at its center was fixed at a height of 52.5 - inch. The top surface of the circular wooden board was painted black as well. For imaging, plants wer e placed under the circular wooden board, and the panicle of interest (primary panicle) was gently passed through the aperture. To adjust for variable plant height, we used an electric scissor lift table (Additional File 2). A metal hook attached to the ceiling of the circular wooden chamber was adhered to top of the panicle for stabilizing the panicle (Additional File 2).

Also, a rotary double-ring apparatus having an inner and an outer ring is fixed on top of the circular wooden board (Additional File 2). A 24-inch aluminum-based outer ring with snow-ball bearings is used to hold two Sony α6500 cameras for imaging and LED lights (ESDDI PLV-380, 15 Watt, 5000 LM, 5600K) for light source, which undergo a 360° rotation around the panicle. The rotation is controlled by an electric motor system. The rotary double-ring apparatus has three major parts: (a) a toothed wheel connected to the electric motor, (b) a small smooth pulley and a cylindrical sleeve used to adjust tension in the belt, and (c) a rotatable ring apparatus that rotates the cameras where the outer ring is covered with a toothed belt. Our camera selection is based on high sensitivity and high stabilization to reduce image distortion during camera motion. The camera also supports customized applications for remote-controlled imaging. We utilized the camera’s time-lapse feature to capture multiple images at the rate of one image per second. Sixty images were captured by each camera per minute, and in total 120 images were taken for each panicle for ea ch time-point and treatment. For labeling, we used quick response (QR) codes as plant identifiers (IDs), which were tagged to the primary panicle. Plant IDs were generated from the images of during the later imaging processing stage. The PI-Plat were constructed mostly using commercial off-the-shelf components at a comparably low cost.

#### Color checkerboards

Color checkerboards printed on white letter-size papers were pasted on all four sides of wooden chamber and on the top surface of the circular wooden board (Additional File 2). Each checkerboard included 20 × 20 squares (1 cm^2^) with colors that were randomly generated in the RGB color space. Color checkerboards were used to provide additional feature points in the 3D reconstruction process. These feature points were used to recover camera parameters, which included the intrinsic calibration (i.e., radial distortion of the lens and the focal length) and the extrinsic calibration (i.e., the position and orientation of the camera) [37]. These additional feature points were important for generating a stable 3D point cloud because the panicle itself had a relatively uniform color and similar patterns, which might not provide enough feature points for the 3D reconstruction without a color checkerboard.

### Image Acquisition

The supplementary video shows image acquisition process using the PI-Plat (Additional File 3). To capture the growth dynamics of panicles, we performed non-destructive imaging of primary panicle corresponding to control and HNT treated plants at one (W1), two (W2) and three-weeks (W3) post-fertilization.

### Image Processing

#### 3D point cloud reconstruction

For 3D point cloud reconstruction, we used the MVE pipeline [37]. First, we converted all the RGB (red, green, and blue) images into the HSV (hue saturation value) space. Then, the background in all images (i.e., the part corresponding to the walls and the circular wooden board) was segmented [43] and removed using the same threshold. With the removal of the background, the amount of feature points in the 3D reconstruction process, as well as the computation time, was reduced. Since all images were taken in the PI-Plat chamber with a constant light, the same threshold worked optimally for all the panicles. Multiple tests using the ‘*colorthresholder*’ application in Matlab showed that the background can be effectively removed if hue, saturation, and value were controlled in the ranges of 0.2-0.5, 0.5-1, and 0.2-0.7, respectively. After background removal, denoising on the images was performed and the components that did not belong to a panicle (e.g., the turntable ring, the residues of checkerboards, etc.) were considered as noise and removed. These pre-processed images were used to reconstruct the 3D point clouds for each panicle at a given time-point. For this, the corresponding feature points in images were detected and matched to form a sparse point cloud in an incremental SfM process. Then, depth maps were reconstructed for each view and merged into a dense point cloud.

#### Trait extraction using 3D point cloud

Once a point cloud at each time point was generated, we were able to extract traits of interest from the reconstructed 3D structure of panicles from these time-varying point clouds. First, each point cloud was segmented into different components (such as a panicle, the color checkboards, and the rotary double-ring apparatus) by leveraging their distinct positions or colors. For example, the color checkboards were approximately located on the boundaries (i.e., the locations of walls and the top surface of the circular wooden board) of a point cloud, and the metal hook was located at the top of the point cloud and has a gray color. Second, the point clouds need to be scaled and aligned, as different point clouds may have different scales and orientations after reconstruction. In this work, the geometries of the color checkboards and the rotary double-ring apparatus were constant during imaging acquisition. Thus, we scaled and aligned the color checkboards and the apparatus across the point clouds. In this way, the rest of the point clouds were scaled and aligned as well, such that panicles in different point clouds can be compared at the same scale [44]. Third, each point cloud was voxelized for volume quantification [45]. The same bounding box was employed to enclose each point cloud. The bounding box was cube-shaped and aligned across the point clouds with respect to the color checkboards and the apparatus. Then, an equivalent discrete voxel-based grid was generated. The grid size was obtained by dividing each edge of the bounding box by 1000. Thus, a volume with a resolution of 1000 × 1000 × 1000 was generated to sample the 3D space. Finally, the points not belonging to a panicle were removed. Therefore, some voxels were filled with a group of panicle points and the other voxels were empty. For each filled voxel, we computed the average color (i.e., RGB) intensity of the points contained in the voxel. Subsequently, the following features were extracted from a volume: (a) voxel count: the number of the filled voxels, and (b) color intensity: the sum of color intensities of all filled voxels.

#### 2D pixel count extraction from multi-view images of developing panicles

For a comparison purpose, conventional 2D based image analysis of panicles was also employed. Specifically, the total pixel count of a panicle was calculated from its corresponding 120 images captured from multiple views. To achieve this, first, each pre-processed image was segmented using the ‘*colorthresholder*’ application in Matlab. This resulted in a set of separated regions. Second, because the checkerboards used in our experiment had green squares whose color was similar to a panicle, the square-shaped regions were detected using solidity. For each region, its solidity is defined as the ratio of the region’s area to the region’s convex hull area. The solidity of each region was calculated using the ‘*regionprops*’ function in Matlab. We did not account for regions that had solidity values larger than 0.7. In addition, given the relatively marginal size of a panicle, a region larger than certain pixels (800 pixels in our study) was filtered out. Therefore, only the pixels of the panicle remained, and the pixel count of the panicle in an image was calculated. We summed the pixel count obtained from each of the 120 multi-view images of the panicle as the total pixel count.

### Scanning of Mature Panicles using Flatbed Scanner

Next, we analyzed mature primary panicle to gain ground truth and derive features, which were compared with the developing panicle. For this, the primary panicles were harvested, and scanned images were obtained using an Epson Expression 12000 XL scanner (600 dpi resolution). Branches on primary panicles were carefully spread out to avoid overlaps in the scanned images. These scanned images were used to extract the following traits: projected surface area of the primary panicle, projected seed count of the primary panicle, average of major (seed length) and minor (seed width) axis, and area of the individual seed on the primary panicle. In this set of images, the panicles were placed over black background. We segmented the panicles from the background using color thresholding and obtained the binary images. As a panicle was mostly yellowish in color and the background was black, an image was transformed in the HSV color space to segment the panicle (setting for range: hue 0-0.3, saturation 0.2-1, and value 0.5-1). In principle, a harvested mature panicle has all the seeds attached to the rachis. Therefore, we first used morphological opening [46] to process the images. As the branches were relatively thin and the seeds were relatively thick, most regions of the seeds were disconnected from each other after morphological opening by removing the branch pixels. As the seeds have an oval shape, the regions that were too thin were removed. The remaining regions corresponded to seeds. The length, width, and area of a seed was calculated from its region using the ‘*regionprops*’ function in Matlab.

### Manual Phenotyping of the Mature Panicle

Next, we manually measured the yield traits on mature primary panicle after harvesting. For this, we collected data for (a) total seed weight, (b) total seed number, (c) weight per seed, and (d) number of fertile and sterile seeds to calculate percentage fertility.

### Correlation Analysis

For pairwise correlation analysis, the 3D reconstruction-based features (voxel count and color intensity) and the total pixel count (2D) derived from the multi-view images of developing panicle were compared with end-point measurements at maturity. For the end-point measurements, the traits derived from flatbed scanned images as well as manual measurements from the primary panicle at maturity were considered. These traits were collected from 11 rice genotypes with two to three replicates per genotype and per treatment (control and HNT). A total of 55 observations were used for Pearson correlation analysis. The correlation analysis was performed using R v. 3.4.3 [47] and RStudio v.1.1.419 [48]. Correlation matrices containing Pearson correlation coefficients and p-values were obtained using the `rcorr` function in “Hmisc” package [49]. Matrix displaying correlation between selected traits was plotted using ‘chart.Correlation’ in the “PerformanceAnalytics” package [50]. Both the raw data and the complete correlation matrix are provided (Additional File 4 and 5).

### Data Accessibility

The text-based raw data generated from 3D reconstruction-based approach, flatbed scanner, and manual measurements for this work is provided as additional files with this submission. Raw image data is large and hence only part of them is shared for user testing on a UNL Box repository (https://unl.box.com/s/g0bof1mpfp33hn66b2qabrk9kiwmhbzv).

## Results

### Workflow of PI-Plat

Evaluation of inflorescence-related parameters is limited by traditional phenotyping methods. Advances in plant phenotyping methodology have enhanced our understanding of vegetative organs and overall plant structures. However, we still need to capitalize on the technological advancement in optics, computer vision, and software design, to capture complex plant structures. In this study, we developed a *P*anicle *I*maging *Plat*form (PI-Plat) to understand yield-related parameters by reconstructing 3D space to derive digital traits (Additional File 2).

For method validation, we used 11 rice genotypes, from the *indica* and *japonica* rice sub-populations (Additional File 1). Once 50% of primary panicle underwent flowering, a subset of plants was maintained under control conditions and the rest were moved to a greenhouse with high night temperature (HNT) condition [51]. The motivation for HNT treatment is to explore the phenotypic variation in rice germplasm as rice grain development is known to be sensitive to HNT [52–54]. The primary panicles from each plant and treatment were imaged three times on a weekly basis (week 1, 2, and 3) using the PI-Plat. For imaging, two visible cameras, held at two different positions, were employed on a rotating imaging system. Sixty images per camera, corresponding to an image clicked every six degrees, aided in capturing multiple views covering 360° of the panicles (Additional File 3). In total, 19,800 images were captured for the 11 genotypes. Each panicle image was segmented and used to reconstruct 3D point clouds which were used to extract phenotypic traits such as (i) voxel count and (ii) color intensity (Figure 1 and Table 1).

**Table 1:**
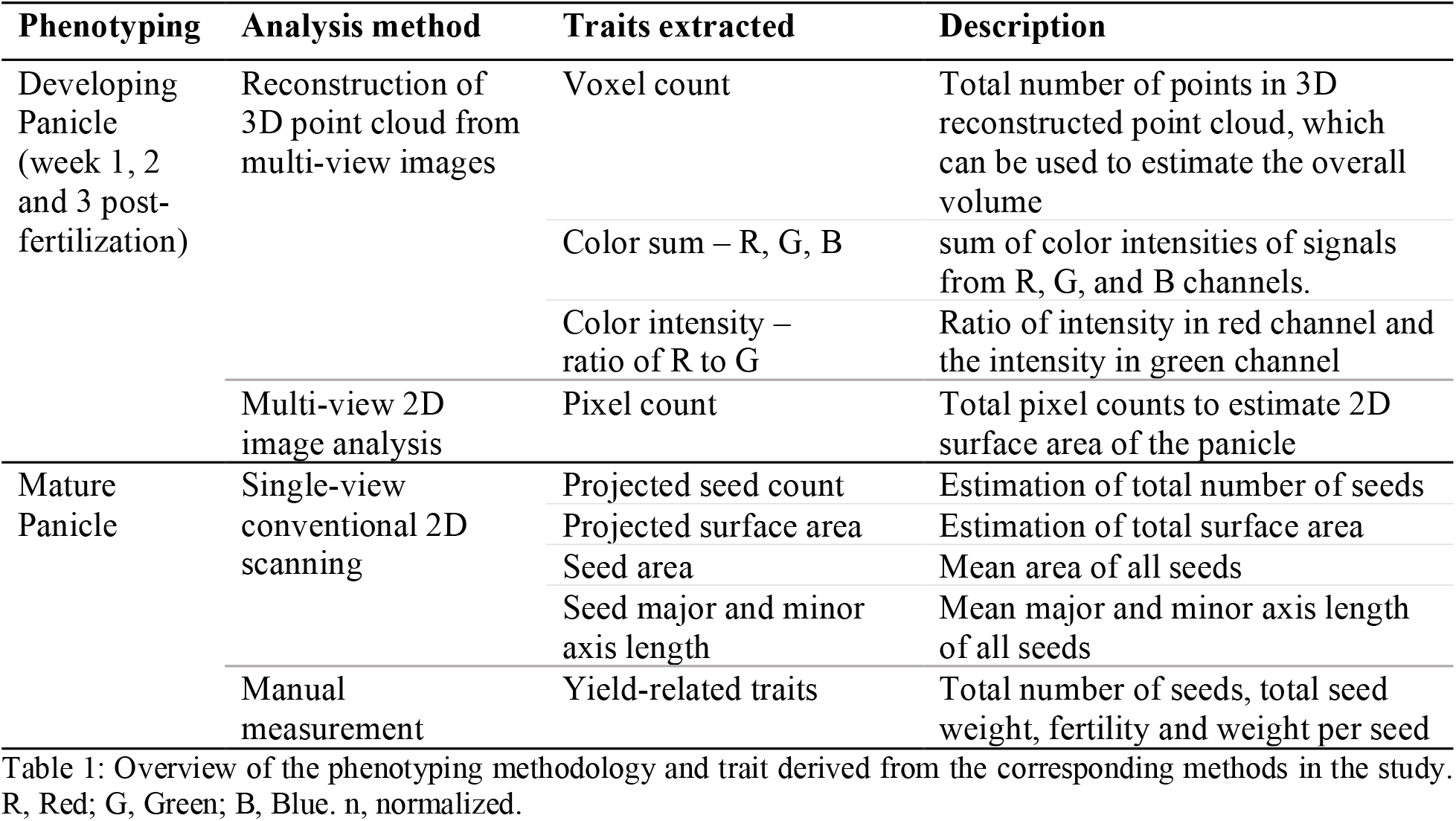
Overview of the phenotyping methodology and trait derived from the corresponding methods in the study. R, Red; G, Green; B, Blue. n, normalized.

### Correlation between traits derived from multi-view images of developing panicle and yield related components at maturity

First, we aimed to determine if the traits derived from 3D reconstruction of the developing panicle correlate with the yield related components at maturity. For this, the 3D reconstruction-based point cloud features derived from multi-view images (voxel count, color intensity) were compared to end-point measurements of the mature panicle (Additional File 5). The end-point measurements correspond to (i) flatbed scanned images (projected surface area at the panicle level, projected seed count, and morphometric measurements at individual seed level; seed area, seed length and width) and (ii) manual measurements (total seed weight, seed number, weight per seed, and fertility) of the mature panicle. Among all the traits derived from 3D reconstruction, only voxel count of developing panicle exhibited significant positive correlation with projected surface area (r_w1_, r_w2_, r_w3_; 0.64, 0.55, 0.82), total seed weight (r_w1_, r_w2_, r_w3_; 0.48, 0.50, 0.74) and seed number (r_w1_, r_w2_, r_w3_; 0.67, 0.61, 0.70) at maturity (Figure 2, Additional File 5). The correlation of the voxel count with projected surface area (r_w1_= 0.64) and total seed weight was relatively low at week 1 (r_w1_: 0.48) and increased with later weeks, week 2 and 3 (r_w1_ < r_w2_ < r_w3_; Figure 2). On the other hand, the correlation between the voxel count of a developing panicle and the seed number at maturity remained stable (Figure 2). Notably, the color intensity derived from 3D reconstruction did not exhibit meaningful correlation with any of the endpoint measurements (Additional File 5).

**Figure 2:**
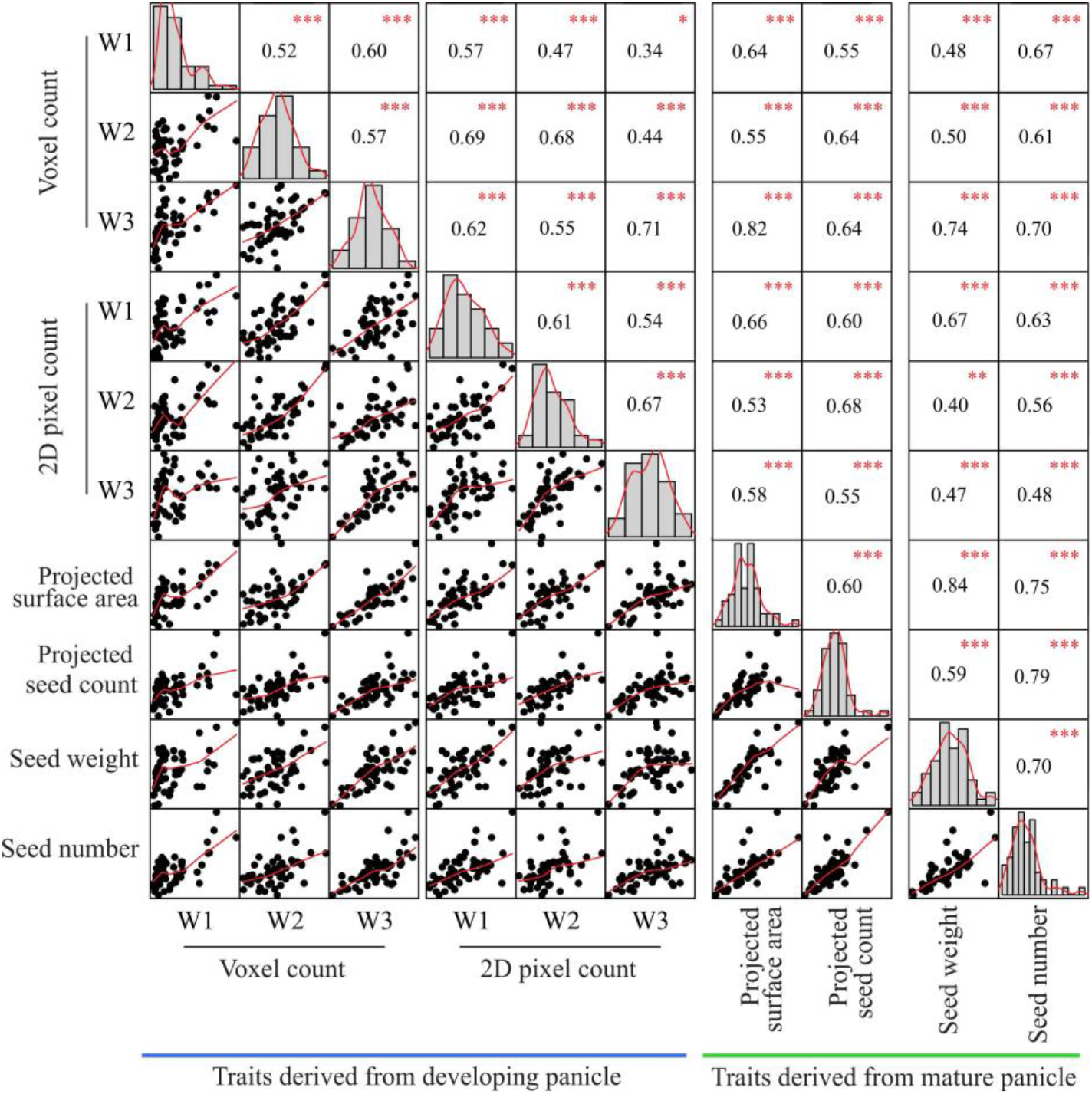
Correlation of traits derived from 3D reconstruction, 2D scanning and manual measurements of inflorescence-related traits. Using PI-Plat, developing panicles were imaged on weekly basis (week 1, 2, and 3). For a respective panicle, multi-view images were used for 3D reconstruction to extract voxel count. Also, 2D pixel count was estimated for developing panicle. Phenotypic traits from mature panicle were analyzed by flatbed scanner (projected surface area and seed count), and manual measurements (seed number and weight). Pearson correlation analysis for traits of primary interest is represented. Similar analysis for other extracted traits is listed in Additional File 4. Histograms and red line represent the distribution of each trait. p-value for significant correlation is shown in red (*** p < 0.001, ** p < 0.01, * p < 0.1), *n* = 55.

Next, the multi-view images were also used to perform the conventional 2D image analysis to extract the total pixel count of a developing panicle for week 1, 2 and 3 (Figure 1). Then, the derived traits at each week were compared with the end-point measurements (Additional File 5). Consequently, the total pixel count showed a positive correlation with all the traits derived from flatbed scanned images and manual measurements at maturity. The correlation between the total pixel count and the projected surface area as well as the total seed weight was unstable. Surprisingly, these correlations at week 3 were lower than the correlations at week 1 (Figure 2).

### Voxel count – an estimate for grain-filling rate

Grain filling rate is the major determinant of mature crop yield. However, evaluating seed weight dynamics usually requires destructive phenotyping methods. In our study, we estimated voxel count from the 3D reconstruction of developing panicles, which represents the overall volume of a panicle, and thus accounts for grain-filling rate. In general, we observed a temporal trend of progressive increase in voxel count over three weeks during the post-fertilization period (Figure 3A). Under control conditions, voxel counts at W2 and W3 were significantly higher than the one at W1, while no significant difference was observed between W2 and W3 (Figure 3A). These results indicate that substantial gain in overall seed volume occurs before W2. Interestingly, plants treated with HNT, possessed significantly higher voxel count at W1 compared to control. These differences dissipated at W2 and W3, as no significant differences between control and HNT treated plants were observed (Figure 3A).

**Figure 3:**
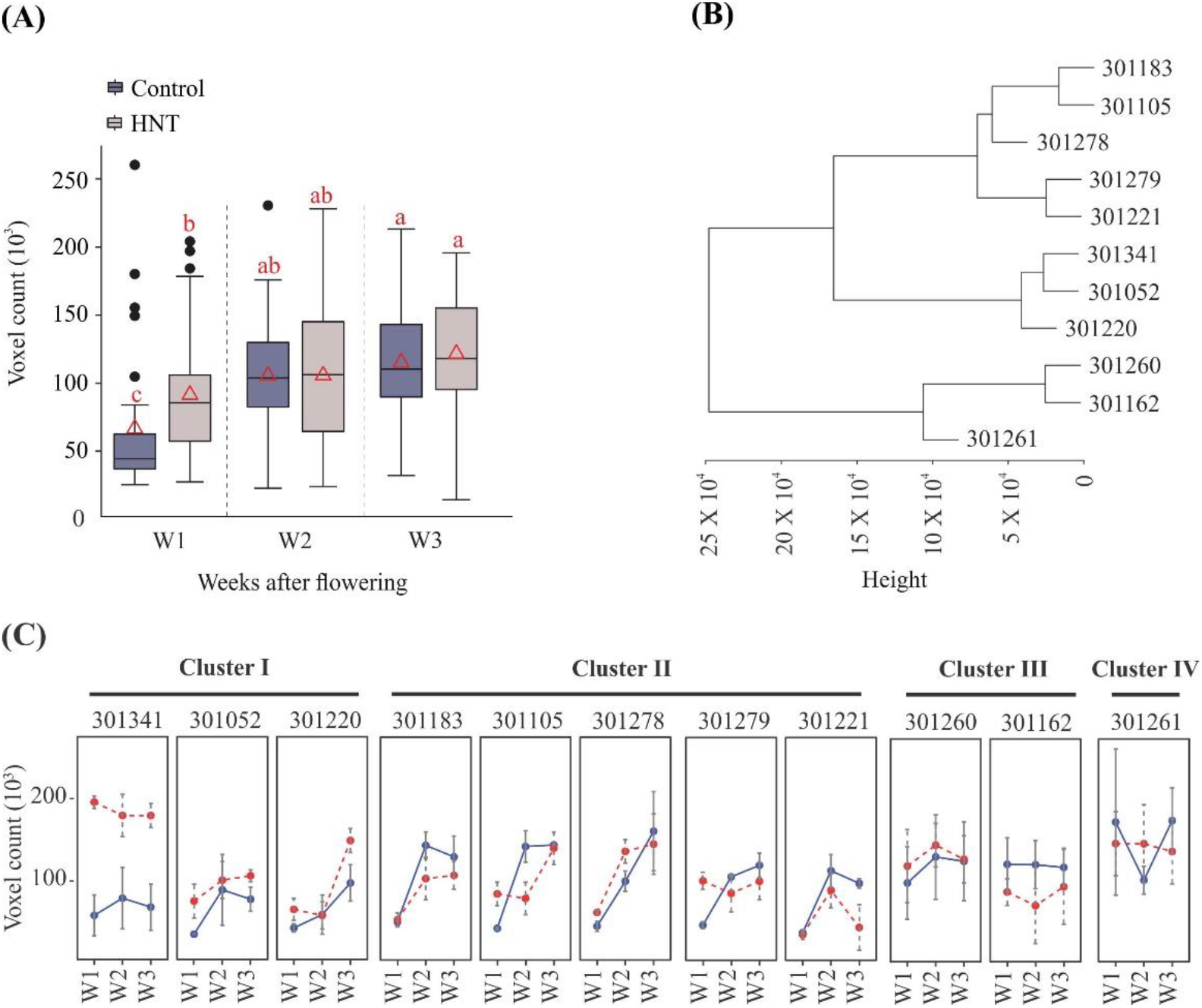
Estimation of voxel count. Voxel count derived from 3D point cloud represents overall volume of developing panicle. **(A)** Average voxel counts from all genotypes for a respective treatment (control and HNT) and time-point (week 1, 2, and 3) is shown. Box plot represents range, median and mean (red triangle) for the same. Means connected with similar letter are not significantly different from each other (Student’s t-test; p < 0.1). (**B**) Hierarchical clustering analysis of genotypes based on their voxel count in control conditions. **(C)** Voxel count for individual genotypes corresponding to cluster I-IV. Y-axis represent voxel count, x-axis indicate time-point (week 1, 2, and 3). C: control, HNT: high night temperature. Box plot represents range, median and mean (red triangle) for the same. Means connected with similar letter are not significantly different from each other (Student’s t-test; p < 0.1).

Next, we evaluated the weekly voxel count for individual genotypes grown under control and HNT stress conditions (Figure 3B and C). We performed hierarchical clustering based on voxel count for control condition panicles (Figure 3B). The analyses grouped 11 genotypes into four distinct clusters (Figure 3B and D).). Cluster I was comprised of 301341, 301052, and 301220, cluster II: 301183, 301105, 301278, 301279, and 301221, cluster III: 301260 and 301262, and, while cluster IV constituted only one genotype, 301261 (Figure 3C). Interestingly, the 4/5 genotypes in Cluster II (301183, 301105, 301221, 301279) showed a significant gain in voxel count between W1 and W2 (Figure 3C). For genotypes in Clusters I, III, and IV, the voxel count trend did not show any significant difference between W1, W2 and W3 (Figure 3C).). This could be because these genotypes may have already gained their potential seed size by W1, and thereby only incremental changes occur afterwards.

### Color intensity – an estimate for rate of maturation

Rate of panicle maturation is a well-studied trait that directly impacts final yield [55, 56]. Heat stress impacts rice seed development and hence alters the panicle maturation rate [57, 58]. Therefore, evaluating the dynamic of panicle maturation could be potentially useful in determining the dynamic of stress response in rice. However, evaluation of the respective traits is done by conventional phenotyping methods, which are inherently laborious and subjective. To estimate the panicle maturation dynamics, we extracted intensity of the RGB channels from the 3D point cloud. Then, we used the ratio of intensity from R to G channels to estimate the yellowness of developing panicle, which increases as the panicles mature. We observed a temporal trend indicating an increase in the ratio of R to G from W1 to W3 (Figure 4A). This observation is consistent with the progression of panicle maturation as its color changes from green to yellow. Interestingly, the R to G ratio was significantly higher for plants treated with HNT compared to control, suggesting that HNT accelerates the rate of panicle maturation. We next explored the genotypic differences for maturation rate (Figure 4B). We observed consistent increase in the R to G ratio from W1 to W3 under control and HNT (Figure 4B). The R to G ratio for majority of genotypes was significantly higher for HNT treated plants than control (Figure 4B and Additional File 5).

**Figure 4:**
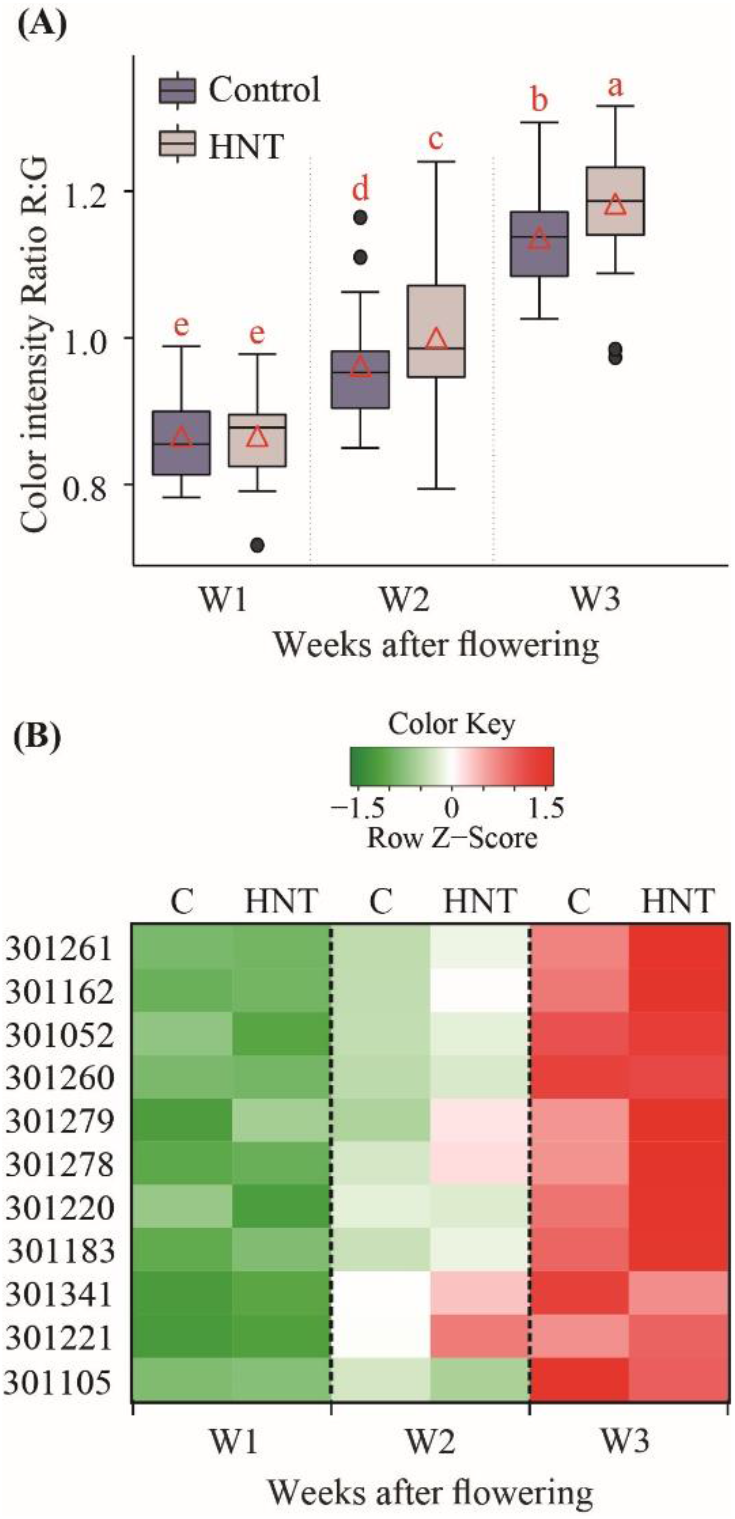
Estimation of color intensity. Color intensity represents sum of color intensities of signals from red (R), green (G), and blue (B) channels. **(A)** Average ratio of R to G intensities from all genotypes for a respective treatment (control or HNT) and time-point (week 1, 2, and 3) is shown. Box plot represents range, media and mean (red triangle) of the R to G ratio. Means connected with same letter are not significantly different from each other (Student’s t - test; p < 0.1). **(B)** Heat map of R to G ratio for different genotypes under control and HNT.

## Discussion

With the recent advances in automated plant image acquisition, accurate quantification of phenotypic traits has become the focal point for realizing the potential of plant phenomics. The primary focus of automated phenotyping platforms has been on the vegetative growth and development and to some extent on the root architectural traits [53–55 and references therein]. Only limited effort has been directed towards more complex yield related traits such as IA in greater detail [16, 20–22, 28, 62]. After flowering, inflorescence undergoes dynamic changes, such as grain filling and maturation, which significantly contributes towards the final yield in cereals. Previous attempts to capture inflorescence-related traits have been limited to end-point measurements. Further, automated Lemnatech phenotyping system, which is mainly used for whole plant imaging, is not suitable to extract high-resolution data from the inflorescence. Hence, the major goal of this study was to capture the growth and developmental dynamics of inflorescence architecture (IA) at high-resolution in rice. To this end, we have developed a low-cost effective system ‘PI-Plat’ to comprehend multi-dimensional features of IA (Figure 1). One of the main novelties of the PI-Plat is that it is designed to reconstruct 3D models of smaller plant parts, in this study ‘panicle’, with a very high resolution. Also, compared to the widely used turntable imaging system where cameras rotate [63], the panicle is fixed at the center of the PI-Plat, thus the vibration is avoided, and the 3D point cloud has less noise. This imaging system can be used to image the panicles in a non-destructive manner, which provides an opportunity to perform temporal phenotyping of the same panicle at consequent developmental stages. On similar basis, rice developing panicles were imaged on weekly basis after fertilization to capture growth dynamics. The multi-view images of developing rice panicle were used for 3D reconstruction, which enabled us to capture digital traits, such as voxel count and color intensity.

We found that the 3D reconstruction-based feature – voxel count has a positive correlation with seed number and total weight at maturity. Panicle development after fertilization involves change in seed weight and volume, but seed number remains constant. Consequently, we observed the temporal trend for correlation of voxel count with final seed weight but not with seed number (Figure 2). Our correlation analysis signifies that image-based phenotyping of developing panicles can be used to estimate the final yield outcome. This information can be valuable for elucidating the physiological and genetic basis of yield components in rice. Various yield components are determined by numerous genes and pathways, which likely influence the yield traits at different developmental phases during panicle development. By using the 3D reconstruction-based voxel count during the panicle development, researchers can identify phenotypic variation over time for divergent genotypes, hence increase the mapping resolution for linking genotypes-to-phenotype. Furthermore, relatively stable correlation between voxel count and seed number at maturity suggest that image-based phenotyping after fertilization can be used to estimate final seed number. In contrast, the 2D based total pixel count of developing panicle showed relatively lower and unstable correlation with seed number and total seed weight at maturity (Figure 2). Interestingly at W3, 2D based pixel counts had lower correlation with endpoint measurements than voxel counts. For instance, the correlation of voxel count with projected surface area and total seed weight was 0.82 and 0.74, respectively, while the correlation of 2D pixel count with projected surface area and total seed weight was 0.58 and 0.47, respectively. This could be due to the limitation of using convention 2D-based phenotyping to completely capture the growth and color dynamics of developing rice seed. Since voxel count positively correlates with final weight, it can be used to capture the weight or volume dynamics. We observed an increase in voxel count from W1 to W3, which is directly related to the increase in size and volume of developing seeds. In context of panicle development, it accounts for rate of grain-filling. Significant gain in the voxel count was achieved by W2 suggesting that substantial seed volume is attained by week 2 (Figure 3). This observation holds true for 4/11 genotypes, while the other seven genotypes do not show such any significant difference between W1, W2, and W3. One possible explanation could be that these genotypes might have accelerated increase in panicle volume and/or seed weight by W1; thus, exhibiting incremental changes during the subsequent two weeks. We observed higher voxel count for HNT treated plants compared to control plants at W1 (Figure 3A). Surprisingly, these differences dissipated at W2 and W3, and no significant difference was observed at maturity. These results highlight the importance of temporal phenotyping relative to single time point measurements. Thus, an end-point measurement approach is not practical to identify and hence map traits that are not persistent at maturity. Since, rice and most other grain crops such as wheat and maize are generally more sensitive to environmental stresses, such as heat and drought, the approach of capturing dynamic reproductive traits in a non-destructive manner will be valuable for research aimed at improving yield resilience to environmental stresses. Early detection of transitory phenotypes/traits is also valuable for molecular studies. Measurement of color intensities from 3D point cloud aided us in understanding the dynamics of panicle maturation for diverse genotypes. Notably, panicles from HNT treated plants showed significantly higher R: G indicating that HNT plants undergo faster maturation. These traits derived from 3D reconstruction of multi-view images provided a close approximation of structural features of the developing rice panicle.

To harness the full potential of the existing genetic resources, we need to bridge the gap between genotype and phenotype. In this context, high throughput genotyping has been facilitated by development of low-cost sequencing platforms. However, accurate and efficient phenotyping of large-scale populations is a major bottleneck for crop improvement. The emergence of phenotyping platforms specifically targeting inflorescence-related traits promise close approximation of the yield-related parameters. PI-Plat provides an important first step towards achieving higher spatial and temporal resolution in IA phenotyping without destructive sampling. The next step towards achieving high-throughput phenotyping of IA traits is the automation for enabling researchers to develop genotype-to-phenotype linkages. Although, the 3D derived voxel count, and color intensity developed as part of PI-Plat can be used to screen large populations elucidating phenotypic variability in inflorescence-related traits, it is still a laborious task given the lack of automation. In summary, PI-Plat-derived 3D traits fills a significant gap in the plant phenotyping toolbox by providing greater spatial and temporal sensitivity of capturing dynamic inflorescence traits, especially for studying abiotic stress responses during reproductive development.

## Abbreviations

IA: inflorescence architecture
PI-Plat: panicle imaging platform
3D: 3 dimensional
2D: 2 dimensional
HNT: high night temperature
W1/2/3: one/two/three week after flowering
RGB: red green and blue

## Authors’ Contribution

HW, HY, YG, PS, and PP conceived and designed the experiment. PP, JS, BKD, FZ, and TG performed the experiments. FZ and TG performed imaging data analysis. PP and JS analyzed the results and wrote the manuscript. All authors read and approved the manuscript.

## Acknowledgements

We thank Martha Rowe for the help with scanning of mature panicles and seeds.

## Availability of data and material

Due to the relatively large size of the raw data, only part of them is shared on a UNL Box repository (https://unl.box.com/s/g0bof1mpfp33hn66b2qabrk9kiwmhbzv). The raw images used for 3D reconstruction and manual phenotyping dataset used in this study is available from the corresponding author on request.

## Consent for publication

Not applicable.

## Ethics approval and consent to participate

Not applicable.

## Additional Files (Additional data is available online)

Additional File 1: Genetic and geographical information of the rice genotypes used in the study.

Additional File 2: PI-Plat and its components.

Additional File 3: Video showing PI-Plat in motion.

Additional File 4: Raw data collected from developing (imaging derived) and mature (manual measurements) panicles.

Additional File 5: Pearson correlation analysis for all the traits derived from 3D reconstruction and multi-view 2D-pixel count analysis of developing panicle (yellow color coded), and mature panicle derived traits from 2D scanning and manual measurement (green color coded). Significant correlation values (p value < 0.05) are highlighted in red font.

Additional File 6: Average intensities of (A) red and (B) green channels from all genotypes for a respective treatment (control and HNT) and time-point (week 1, 2, and 3).

## References

1. Tester M, Langridge P. Breeding technologies to increase crop production in a changing world. Science (80-). 2010;327:818–22. doi:10.1126/science.1183700.

2. Alexandratos N, Bruinsma J. World Agriculture towards 2030/2050: the 2012 revision. 2012. www.fao.org/economic/esa.

3. Röth S, Paul P, Fragkostefanakis S. Plant heat stress response and thermotolerance. 2016. doi: 10.1007/978-81-322-2662-8_2

4. Ray DK, Mueller ND, West PC, Foley JA. Yield Trends Are Insufficient to Double Global Crop Production by 2050. PLoS One. 2013;8:e66428. doi:10.1371/journal.pone.0066428.

5. Godfray HCJ, Beddington JR, Crute IR, Haddad L, Lawrence D, Muir JF, et al. Food security: the challenge of feeding 9 billion people. Science (80-). 2010;327:812–8. doi:10.1126/science.1185383.

6. Foley JA, Ramankutty N, Brauman KA, Cassidy ES, Gerber JS, Johnston M, et al. Solutions for a cultivated planet. Nature. 2011;478:337–42. doi:10.1038/nature10452.

7. Richards RA. Selectable traits to increase crop photosynthesis and yield of grain crops. J Exp Bot. 2000;51 suppl_1:447–58. doi:10.1093/jexbot/51.suppl_1.447.

8. Evans LT, Fischer RA. Yield Potential: Its defination, measurement, and significance. Crop Sci. 1999;39:1544. doi:10.2135/cropsci1999.3961544x.

9. Doust A. Architectural evolution and its implications for domestication in grasses. Ann Bot. 2007;100:941–50. https://academic.oup.com/aob/article-abstract/100/5/941/135949. Accessed 14 Mar 2019.

10. Duan L, Yang W, Huang C, Liu Q. A novel machine-vision-based facility for the automatic evaluation of yield-related traits in rice. Plant Methods. 2011;7:44. doi:10.1186/1746-4811-7-44.

11. Reuzeau C, Pen J, Frankard V, Wolf J, Peerbolte R, Broekaert W, et al. TraitMill: a discovery engine for identifying yield-enhancement genes in cereals. Plant Gene Trait. 2010;1. http://biopublisher.ca/index.php/pgt/article/html/53.

12. Granier C, Aguirrezabal L, Chenu K, Cookson SJ, Dauzat M, Hamard P, et al. PHENOPSIS, an automated platform for reproducible phenotyping of plant responses to soil water deficit in *Arabidopsis thaliana* permitted the identification of an accession with low sensitivity to soil water deficit. New Phytol. 2006;169:623–35. doi:10.1111/j.1469-8137.2005.01609.x.

13. Golzarian MR, Frick RA, Rajendran K, Berger B, Roy S, Tester M, et al. Accurate inference of shoot biomass from high-throughput images of cereal plants. Plant Methods. 2011;7:2. doi:10.1186/1746-4811-7-2.

14. Yang W, Xu X, Duan L, Luo Q, Chen S, Zeng S, et al. High-throughput measurement of rice tillers using a conveyor equipped with x-ray computed tomography. Rev Sci Instrum. 2011;82:025102. doi:10.1063/1.3531980.

15. Bylesjö M, Segura V, Soolanayakanahally RY, Rae AM, Trygg J, Gustafsson P, et al. LAMINA: a tool for rapid quantification of leaf size and shape parameters. BMC Plant Biol. 2008;8:82. doi:10.1186/1471-2229-8-82.

16. Wilson Z, Greenberg AJ, McCouch SR, Crowell S, Falcao AX, Shah A. High-Resolution Inflorescence Phenotyping Using a Novel Image-Analysis Pipeline, PANorama. Plant Physiol. 2014;165:479–95.

17. Yazdanbakhsh N, Fisahn J. High throughput phenotyping of root growth dynamics, lateral root formation, root architecture and root hair development enabled by PlaRoM. Funct Plant Biol. 2009;36:938. doi:10.1071/FP09167.

18. Wang L, Uilecan I, Assadi A, … CK-P, 2009 U. HYPOTrace: image analysis software for measuring hypocotyl growth and shape demonstrated on Arabidopsis seedlings undergoing photomorphogenesis. Plant Physiol. 2009;149:1632–7. http://www.plantphysiol.org/content/149/4/1632.short.

19. Fiorani F, Schurr U. Future Scenarios for Plant Phenotyping. Annu Rev Plant Biol. 2013;64:267–91. doi:10.1146/annurev-arplant-050312-120137.

20. Ikeda M, Hirose Y, Takashi T, Shibata Y, Yamamura T, Komura T, et al. Analysis of rice panicle traits and detection of QTLs using an image analyzing method. Breed Sci. 2010;:55–64. https://www.jstage.jst.go.jp/article/jsbbs/60/1/60_1_55/_article/-char/ja/. Accessed 12 Mar 2019.

21. AL-Tam F, Adam H, Anjos A, Lorieux M, Larmande P, Ghesquière A, et al. P-TRAP: a Panicle Trait Phenotyping tool. BMC Plant Biol. 2013;13:122. doi:10.1186/1471-2229-13-122.

22. Zhou Y, Srinivasan S, Mirnezami SV, Kusmec A, Fu Q, Attigala L, et al. Semiautomated Feature Extraction from RGB Images for Sorghum Panicle Architecture GWAS. Plant Physiol. 2018;179:24–37.

23. Aquino A, Millan B, Gaston D, Diago M-P, Tardaguila J, Aquino A, et al. vitisFlower®: Development and Testing of a Novel Android-Smartphone Application for Assessing the Number of Grapevine Flowers per Inflorescence Using Artificial Vision Techniques. Sensors. 2015;15:21204–18. doi:10.3390/s150921204.

24. Millan B, Aquino A, Diago MP, Tardaguila J. Image analysis-based modelling for flower number estimation in grapevine. J Sci Food Agric. 2017;97:784–92. doi:10.1002/jsfa.7797.

25. Wang Z, Underwood J, Walsh KB. Machine vision assessment of mango orchard flowering. Comput Electron Agric. 2018;151:501–11. doi:10.1016/J.COMPAG.2018.06.040.

26. Ji W, Zhao D, Cheng F, Xu B, Zhang Y. Automatic recognition vision system guided for apple harvesting robot. Comput Electr Eng. 2012;38:1186–95. https://www.sciencedirect.com/science/article/pii/S0045790611001819.

27. Crowell S, Falcão A, Shah A, Wilson Z, Greenberg AJ, McCouch S. High-resolution inflorescence phenotyping using a novel image-analysis pipeline, PANorama. Plant Physiol. 2014;165:479–95. http://www.plantphysiol.org/content/165/2/479.short.

28. Gage JL, Miller ND, Spalding EP, Kaeppler SM, De Leon N. TIPS: a system for automated image-based phenotyping of maize tassels. Plant Methods. 2017;13. doi:10.1186/s13007-017-0172-8.

29. Nhung KT, Giang Khong N, Loan TH, Thu NGUYEN D, Chung MAI D, Giang HOANG T, et al. A genome-wide association study using a Vietnamese landrace panel of rice (Oryza sativa) reveals new QTLs controlling panicle morphological traits. BMC Plant Biol. 2018;18. doi:10.1186/s12870-018-1504-1.

30. Adriani DE, Dingkuhn M, Dardou A, Adam H, Luquet D, Lafarge T. Rice panicle plasticity in Near Isogenic Lines carrying a QTL for larger panicle is genotype and environment dependent. Rice. 2016;9:28. doi:10.1186/s12284-016-0101-x.

31. Rebolledo MC, Peña AL, Duitama J, Cruz DF, Dingkuhn M, Grenier C, et al. Combining Image Analysis, Genome Wide Association Studies and Different Field Trials to Reveal Stable Genetic Regions Related to Panicle Architecture and the Number of Spikelets per Panicle in Rice. Front Plant Sci. 2016;7:1384. doi:10.3389/fpls.2016.01384.

32. Li D, Xu L, Tang XS, Sun S, Cai X, Zhang P. 3D imaging of greenhouse plants with an inexpensive binocular stereo vision system. Remote Sens. 2017;9.

33. Omasa K, Hosoi F, Botany AK-J of experimental, 2006 U. 3D lidar imaging for detecting and understanding plant responses and canopy structure. J Exp Bot. 2007;58:881–98. https://academic.oup.com/jxb/article-abstract/58/4/881/425236. Accessed 15 Mar 2019.

34. McCormick RF, Truong SK, Mullet JE. 3D Sorghum Reconstructions from Depth Images Identify QTL Regulating Shoot Architecture. Plant Physiol. 2016;172:823–34. doi:10.1104/pp.16.00948.

35. Brooks MJ, de Agapito L, Huynh DQ, Baumela L. Towards robust metric reconstruction via a dynamic uncalibrated stereo head. Image Vis Comput. 1998;16:989–1002. doi:10.1016/S0262-8856(98)00064-X.

36. Negahdaripour S, Hayashi BY, Aloimonos Y. Direct motion stereo for passive navigation. IEEE Trans Robot Autom. 1995;11:829–43. doi:10.1109/70.478430.

37. Fuhrmann S, Langguth F, Goesele M. MVE – A Multi-View Reconstruction Environment. EUROGRAPHICS Work Graph Cult Herit. 2014.

38. Sodhi P, Vijayarangan S, Wettergreen D. In-field segmentation and identification of plant structures using 3D imaging. In: 2017 IEEE/RSJ International Conference on Intelligent Robots and Systems (IROS). IEEE; 2017. p. 5180–7. doi:10.1109/IROS.2017.8206407.

39. Vijayarangan S, Sodhi P, Kini P, Bourne J, Du S, Sun H, et al. High-Throughput Robotic Phenotyping of Energy Sorghum Crops. In: n: Hutter M., Siegwart R. (eds) Field and Service Robotics. Springer Proceedings in Advanced Robotics, vol 5. Springer, Cham. Springer, Cham; 2018. p. 99–113. doi:10.1007/978-3-319-67361-5_7.

40. Duan T, Chapman SC, Holland E, Rebetzke GJ, Guo Y, Zheng B. Dynamic quantification of canopy structure to characterize early plant vigour in wheat genotypes. J Exp Bot. 2016;67:4523–34. doi:10.1093/jxb/erw227.

41. Hartmann A, Czauderna T, Hoffmann R, Stein N, Schreiber F. HTPheno: An image analysis pipeline for high-throughput plant phenotyping. BMC Bioinformatics. 2011;12:148. doi:10.1186/1471-2105-12-148.

42. Chaudhury A, Barron JL. Machine Vision System for 3D Plant Phenotyping. IEEE/ACM Trans Comput Biol Bioinforma. 2018;:1–1. doi:10.1109/TCBB.2018.2824814.

43. Vantaram S, Saber E. Survey of contemporary trends in color image segmentation. J Electron Imaging. 2012;21. https://www.spiedigitallibrary.org/journals/Journal-of-Electronic-Imaging/volume-21/issue-4/040901/Survey-of-contemporary-trends-in-color-image-segmentation/10.1117/1.JEI.21.4.040901.short.

44. Besl P, McKay N. Method for registration of 3-D shapes. Sens Fusion IV Control Paradig. 1992. https://www.spiedigitallibrary.org/conference-proceedings-of-spie/1611/0000/Method-for-registration-of-3-D-shapes/10.1117/12.57955.short.

45. Cohen-Or D, Kaufman A. Fundamentals of surface voxelization. Graph Model image Process. 1995. https://www.sciencedirect.com/science/article/pii/S1077316985710398. Accessed 8 Apr 2019.

46. Gonzalez R, Woods R. Digital Image Processing, Global Edition. 2018.

47. R Core Team. R: A Language and Environment for Statistical Computing. 2017.

48. RStudio Team. RStudio: Integrated Development Environment for R. 2016.

49. Frank E, Harrell J, with contributions from Charles Dupont and many others. Hmisc: Harrell Miscellaneous. R package version 4.1-1. 2018.

50. Peterson BG, Peter C. PerformanceAnalytics: Econometric Tools for Performance and Risk Analysis. R package version 1.5.2. 2018.

51. Dhatt BK, Abshire N, Paul P, Hasanthika K, Sandhu J, Zhang Q, et al. Metabolic dynamics of developing rice seeds under high night-time temperature stress. Front Plant Sci. 2019;10:1443.

52. Peng S, Huang J, Sheehy JE, Laza RC, Visperas RM, Zhong X, et al. Rice yields decline with higher night temperature from global warming. PNAS. 2004;101:9971–5. www.pnas.orgcgidoi10.1073pnas.0403720101.

53. Cheng W, Sakai H, Yagi K, Hasegawa T. Interactions of elevated [CO2] and night temperature on rice growth and yield. Agric For Meteorol. 2009;149:51–8. doi:10.1016/J.AGRFORMET.2008.07.006.

54. Coast O, Ellis RH, Murdoch AJ, Quiñones C, Jagadish KS V. High night temperature induces contrasting responses for spikelet fertility, spikelet tissue temperature, flowering characteristics and grain quality in rice. Funct Plant Biol. 2015;42:149. doi:10.1071/FP14104.

55. Jongkaewwattana S, Geng S, Hill JE, Miller BC. Within-Panicle Variability of Grain Filling in Rice Cultivars with Different Maturities. J Agron Crop Sci. 1993;171:236–42. doi:10.1111/j.1439-037X.1993.tb00135.x.

56. Ellis RH. Rice seed quality development and temperature during late development and maturation. Seed Sci Res. 2011;21:95–101. doi:10.1017/S0960258510000425.

57. Begcy K, Sandhu J, Walia H. Transient Heat Stress During Early Seed Development Primes Germination and Seedling Establishment in Rice. Front Plant Sci. 2018;9:1768.

58. Folsom JJ, Begcy K, Hao X, Wang D, Walia H. Rice fertilization-Independent Endosperm1 regulates seed size under heat stress by controlling early endosperm development. Plant Physiol. 2014;165:238–48. doi:10.1104/pp.113.232413.

59. Humplík JF, Lazár D, Husičková A, Spíchal L. Automated phenotyping of plant shoots using imaging methods for analysis of plant stress responses – a review. Plant Methods. 2015;11:29. doi:10.1186/s13007-015-0072-8.

60. Li L, Zhang Q, Huang D, Li L, Zhang Q, Huang D. A Review of Imaging Techniques for Plant Phenotyping. Sensors. 2014;14:20078–111. doi:10.3390/s141120078.

61. Fahlgren N, Gehan MA, Baxter I. Lights, camera, action: high-throughput plant phenotyping is ready for a close-up. Curr Opin Plant Biol. 2015;24:93–9. doi:10.1016/J.PBI.2015.02.006.

62. Xiong X, Duan L, Liu L, Tu H, Yang P, Wu D, et al. Panicle-SEG: A robust image segmentation method for rice panicles in the field based on deep learning and superpixel optimization. Plant Methods. 2017;13:1–15. doi:10.1186/s13007-017-0254-7.

63. He JQ, Harrison RJ, Li B. A novel 3D imaging system for strawberry phenotyping. Plant Methods. 2017;13:1–8.

