## Supplementary figures and images for "PI-Plat: A high-resolution image-based 3D reconstruction method to estimate growth dynamics of rice inflorescence traits"

### Additional File 2.jpg

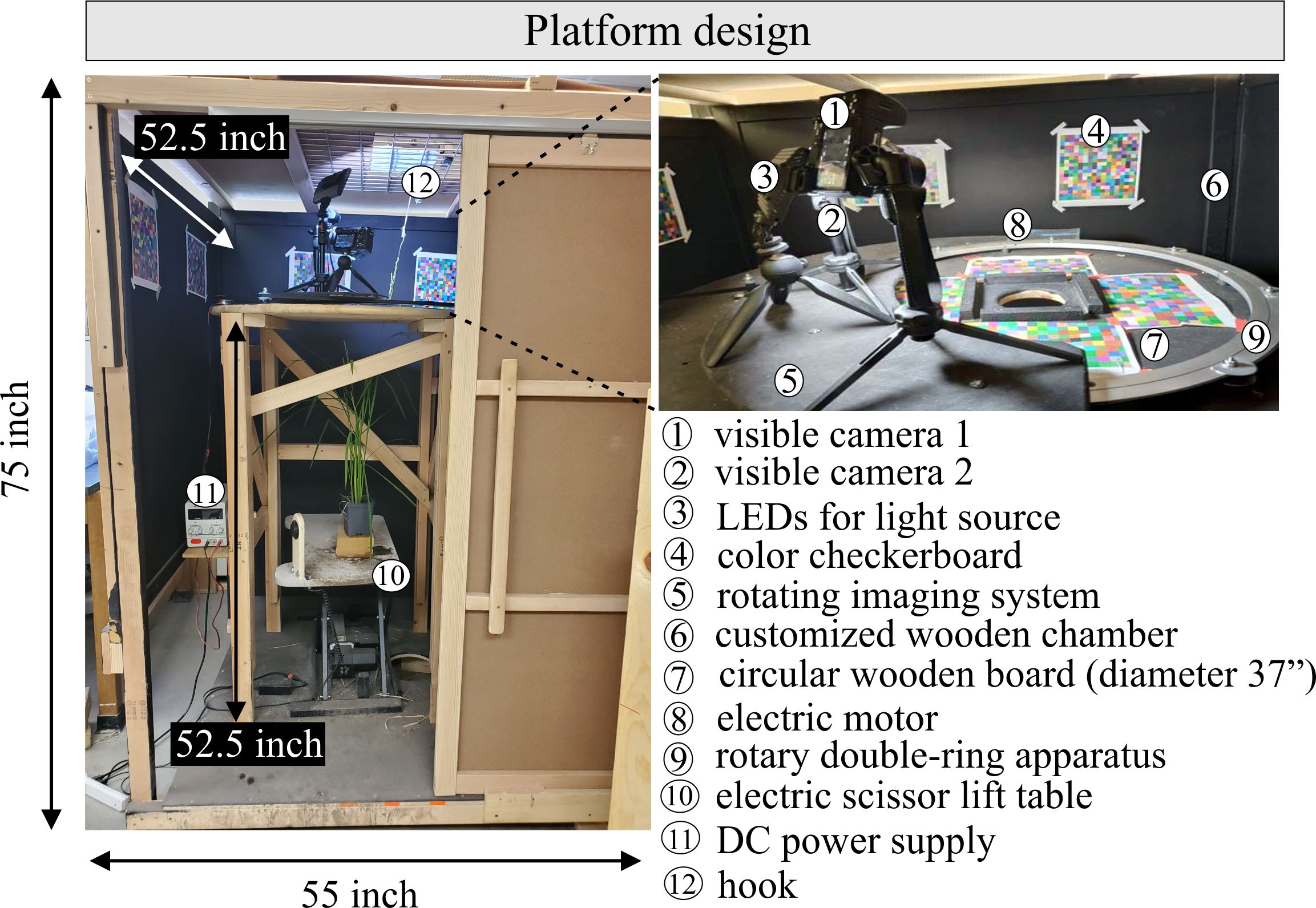

### Additional File 6.jpg

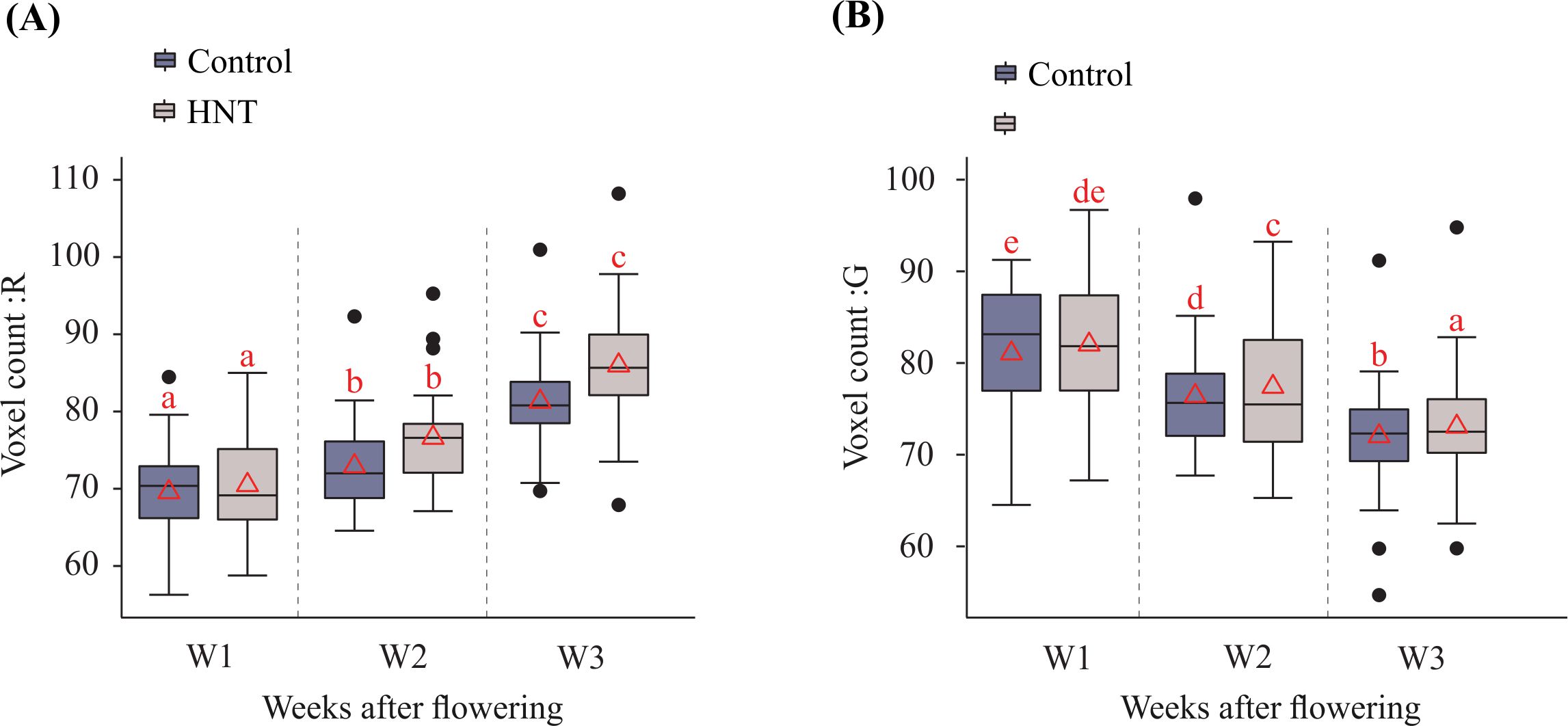
